# Developing hippocampal spheroids model ictogenesis and epileptogenesis

**DOI:** 10.1101/2023.01.06.523024

**Authors:** John Wesley Ephraim, Davide Caron, Angel Canal-Alonso, Juan Manuel Corchado, Gemma Palazzolo, Gabriella Panuccio

## Abstract

Three-dimensional (3D) neural cell cultures inherently lend themselves to high-throughput network electrophysiology studies addressing brain function in health and disease in a more realistic architectural complexity than two-dimensional neural networks. Epilepsy is the emblem of brain network disorders, as it reflects aberrant circuit reorganization and hyper-synchronization, resulting in sudden and uncontrolled electrical discharges (seizures). Modeling the features of epilepsy has so far relied on pharmacological, ionic or genetic manipulation of cells, ex-vivo brain tissue or intact animals, failing to recapitulate most of the epilepsies, which are triggered by unknown causes. Here, we report the spontaneous emergence of epileptiform patterns in spheroids of rodent primary hippocampal cells cultured in physiological condition, i.e., in the absence of a known initiating insult, detected by microelectrode array electrophysiology. Three distinct electrical phenotypes, i.e. interictal (between seizures), ictal (seizure) or mixed, arise from DIV10 to DIV35. In particular, the tonic-clonic ictal discharges become the most prominent at DIV28-35. These patterns exhibit electrographic and spectral features that strikingly resemble those observed in the hippocampus of in vitro and in vivo rodent epilepsy models, as well as of drug-resistant epileptic humans. Remarkably, not all spheroids exhibit full-blown ictal activity, bringing parallelism with the yet unanswered question of why a brain becomes epileptic and a seizure is generated. This evidence warrants caution against hippocampal cell-based therapies for regenerative purposes, as they may initiate epileptogenesis; at the same time, hippocampal spheroids lend themselves as reductionist model supporting high-throughput pre-clinical research on epileptic syndromes involving the hippocampus.

## Introduction

The hippocampus is a highly excitable region within the limbic system, where it governs cognitive processes such as learning and memory via its highly synchronized electrical patterns. Because of its high excitability and rhythmic operating mode, the hippocampus is prone to generate seizures under permissive conditions that impair its finely tuned input/output function. In particular, functional and structural alterations of the hippocampus are the hallmark of mesial temporal lobe epilepsy (MTLE)^1^, the most frequent epileptic syndrome in adults and the most often refractory to medical therapy^2^.

The clinical history of MTLE patients is characterized by a brain insult during childhood, followed by a latent period during which the brain undergoes cellular and network rearrangements that eventually lead to chronic MTLE in adulthood. Established animal models^3^ have enabled a remarkable advancement of our understanding of epileptogenesis and ictogenesis; however, they are bound to a specific initiating condition which may not recapitulate realistically the clinical history of MTLE patients; further, model-to-model and intra-model variability challenge basic and translational research. As a result, we are still unable to answer why a brain becomes epileptic, a seizure is generated, and the patient stops responding to the anti-epileptic drug treatment. In turn, this knowledge gap affects the efficacy of available pharmacological and brain stimulation therapies, which remain symptomatic treatments rather than being aimed at healing the condition. Most importantly, we are to date unable to identify patients at risk of developing epilepsy so to implement a timely preventive therapy.

The lack of effective strategies to prevent or treat MTLE and epilepsy patients overall calls for radically new therapeutic approaches than what available today. Regenerative medicine is the latest frontier to address this unmet clinical need, as it promises to heal brain dysfunction by rebuilding compromised brain circuits. However, the quest for effective and safe brain regenerative approaches is still in its infancy. While the advent of human brain organoids along with the advancement of tissue bioengineering techniques represent a landmark toward the future of brain regeneration^4^, their long culturing time and high cost do not comply with the urgent and massive request; in particular, we are still unable to reliably obtain electrically active hippocampal organoids ^5-7^. Further, there is still little evidence about the suitability and safety of bioengineered brain tissue grafts for brain regeneration^8,9^. At variance, early experimental evidence prior to the advent of brain organoids pointed to the possibility of uncontrolled pathological behavior of hippocampal grafts; specifically, it has been shown that rodent fetal hippocampi grafted into the adult rodent brain can become an epileptic focus^10,11^, and that two-dimensional hippocampal microcultures exhibit spontaneous epileptiform patterns stemming from recurrent excitatory connections and autapses^12,13^. On the contrary, grafting of primary or stem cell-derived interneurons^14^ or of hippocampal stem cells^6^ in epilepsy animal models has been shown to improve the disorder. Thus, there is still little and contrasting evidence about the efficacy and safety of cell-based approaches for brain regeneration and repair. This warrants an in-depth large-scale pre-clinical evaluation in animal models^15^.

To shed light on the potential application of hippocampal replicas for regenerative therapy, we have evaluated the electrophysiological features of hippocampal spheroids obtained from primary rat cells and cultured in physiological condition. In performing acute electrophysiology recordings by means of 3D microelectrode array (MEA) at different time points, we have discovered that hippocampal spheroids spontaneously generate epileptiform activity. This discovery warrants caution in devising cell-based regenerative therapies for the brain; on the other hand, it brings a new high-throughput reductionist model for pre-clinical studies of epileptic syndromes involving the hippocampus.

## Materials and Methods

### Ethics

All procedures involving animals have been approved by the Institutional Animal Welfare Body and by the Italian Ministry of Health (authorization 176AA.NTN9), in accordance with the National Legislation (D.Lgs. 26/2014) and the European Directive 2010/63/EU. Animals were purchased from Charles River, Italy. Human hippocampal EEG recordings from patients diagnosed with drug-resistant epilepsy were kindly provided by the Department of Neurology of the University of Bern with the patients’ consent.

### Ex-vivo sample preparation and maintenance

#### Hippocampal spheroids

Primary hippocampal cells were extracted from E17.5 Sprague-Dawley rat embryos as previously described^16^. In detail, the embryos were decapitated upon extraction, their brains were removed and placed in cold HBSS (Gibco, 14170-88). Approximately 7 hippocampi were placed in 5 ml of Trypsin-EDTA (Gibco, 25200-056) and incubated in the water bath at 37°C for 30 min. After the incubation, 5 ml of complete Neurobasal medium were added and centrifuged for 5 min at 335 x g (Beckman Coulter Allegra X-12R centrifuge). The complete Neurobasal medium consisted of: Neurobasal A (Gibco, 10888022), 2% B27 (Gibco, 17504044), 1% PenStrep (Gibco, 15070063), 1% Glutamax (Thermo Scientific, 35050038) + 10% Fetal Bovine Serum (FBS, Gibco, 10270-106). The supernatant resulting from centrifugation was then discarded and replaced with 3-4 ml of fresh Neurobasal complete medium +10% FBS, and the hippocampi were mechanically dissociated by gently pipetting. The hippocampi suspension was then filtered with a cell strainer (40 mm pore size) and centrifuged for 7 min at 114 x g. Following centrifugation, the supernatant was discarded, and the cells were resuspended in complete Neurobasal medium and counted.

For each spheroid, 30,000 cells were seeded in each well of an ultra-low adhesive U-shaped 96-well plate (CellCarrier™ −96, PerkinElmer) with 200 ml of complete Neurobasal medium. On the third day after seeding, the spheroids were transferred to a 24-well plate coated with 3% agarose (UV-treated for 15-20 min), one spheroid per well. Neurobasal medium was replaced weekly by 50%. All the cultures were incubated at 37°C and 5% CO_2_.

#### Brain slices

Horizonal hippocampus-cortex (CTX) slices, 400 m thick, were prepared from male CD1 mice 4-8 weeks old. Animals were decapitated under deep isoflurane anesthesia, their brain was removed within 60 s, and immediately placed into ice-cold (∼2 °C) sucrose-based artificial cerebrospinal fluid (sucrose-ACSF) composed of (mM): Sucrose 208, KCl 2, KH_2_PO_4_ 1.25, MgCl_2_ 5, MgSO_4_, CaCl_2_ 0.5, D-glucose 10, NaHCO_3_ 26, L-Ascorbic Acid 1, Pyruvic Acid 3. The brain was let chill for ∼2 min before slicing in ice-cold sucrose-ACSF using a vibratome (Leica VT1000S, Leica, Germany). Brain slices were immediately transferred to a submerged holding chamber containing room-temperature holding ACSF composed of (mM): NaCl 115, KCl 2, KH_2_PO_4_, 1.25, MgSO_4_ 1.3, CaCl_2_ 2, D-glucose 25, NaHCO_3_ 26, L-Ascorbic Acid 1. After at least 60 minutes of recovery, individual slices were transferred to a submerged incubating chamber containing warm (∼32 °C) holding ACSF for 20-30 minutes (pre-warming) and subsequently incubated for at least 60 in warm ACSF containing the K^+^ channel blocker 4-aminopyridine^17^ (4AP, 250 M), in which MgSO_4_ concentration was lowered to 1 mM (4AP-ACSF, *cf*. ^18^). In all brain slices, the CA3 output to the CTX was prevented by Schaffer Collateral’s disruption to release cortical ictogenicity^19^. All solutions were constantly equilibrated at pH = ∼7.35 with 95% O_2_ / 5% CO_2_ gas mixture (carbogen) and had an osmolality of 300-305 mOsm/Kg. Chemicals were purchased from Sigma-Aldrich.

### Microelectrode array electrophysiology

Extracellular recordings were performed acutely from all samples through the MEA2100-mini-HS60 amplifier using the Multichannel Experimenter software; the amplifier was connected to the IFB v3.0 multiboot interface board through the SCU signal collector unit. The recording temperature was controlled by a TC02 thermostat and checked with a k-type thermocouple in all experiments. The equipment for MEA recording and temperature control was purchased from Multichannel Systems (MCS), Reutlingen, Germany.

#### Hippocampal spheroids

Individual samples were taken from the incubator immediately prior to the recording and were accommodated onto an 8 × 8 3D MEA (TiN electrodes, diameter 12 µm, height 80 µm, inter-electrode distance 200 µm, impedance ∼150 kΩ, internal reference electrode) with default ring recording chamber. To avoid damaging the 3D MEA electrodes or the spheroid, we did not use a holding anchor. In this setting, the electrical coupling of the spheroid with the MEA was still reliably preserved throughout the recording.

Spheroids were let habituate for 20 minutes before recording. Recordings were performed at ∼37° C, achieved with the use of a custom-made heating lid covering the headstage along with the warming of the MEA amplifier base (temperature set at 37° C). The recording ACSF was equilibrated at pH ∼7.4 through humidified carbogen delivered via a tubing connected to the heating lid and consisted of (mM): NaCl 117, KCl 3.75, KH_2_PO_4_, 1.25, MgSO_4_ 0.5, CaCl_2_ 2.5, D-glucose 25, NaHCO_3_ 26, L-Ascorbic Acid 1, pH 7.4. Signals were sampled at 20 kHz and low-pass filtered at 10 kHz before digitization.

#### Brain slices

Individual brain slices were accommodated onto a 6 × 10 planar MEA (TiN electrodes, diameter 30 μm, inter-electrode distance 500 μm, impedance < 100 kΩ), in which a custom-made low-volume (∼500 μl) recording chamber (Crisel Instruments, Italy) replaced the default MEA ring (*cf*. ^18^) to attain a laminar flow. Brain slices were held down by a custom-made stainless steel/nylon mesh anchor and continuously perfused at ∼1 ml/min with 4AP-ACSF, equilibrated with carbogen. As the custom chamber covered the internal reference electrode, this was replaced by an external reference consisting of a saturated KCl pellet. Recordings were performed at 32° C, achieved with the use of a heating canula (PH01) inserted at the recording chamber inlet port (temperature set at 37° C) along with mild warming of the MEA amplifier base (temperature set at 32° C). Signals were sampled at 5 kHz and low-pass filtered at 2 kHz before digitization.

#### EEG recording

##### Epileptic rats

Hippocampal EEG recordings from epileptic rats were taken from baseline recordings described in^20^ and openly available at 10.17605/OSF.IO/UK94M. Detailed methods are available in the original publication.

##### Epileptic human

Hippocampal EEG recordings from patients diagnosed with drug-resistant epilepsy were acquired with a BrainScope system (M&I, BrainScope, Czech Republic) at a sampling frequency of 25 kHz. The depth electrode arrays were 5, 10 and 15 contact semi-flexible platinum electrodes (ALCIS - Temis Health, France), with a diameter of 0.8 mm, a contact length of 2 mm, a contact surface area of 5.02 mm^2^ and an inter-contact distance of 1.5 mm.

### Image analysis

Phase contrast images were acquired via a Nikon Eclipse Ts2 inverted phase contrast microscope using a 4X objective lens. Images were analyzed using the open-source software Fiji. In detail, the maximum diameter [mm] was taken as the maximum value measured by manually drawing it across different directions. The surface area was measured semi-automatically: to this end, we first set the threshold for automatic detection of the spheroid edges and then used the functions *Set Measurements* and *Measure* for automatic area size calculation.

### Signal processing

MEA signals from brain slices and spheroids, and rodent EEG signals from rodents were automatically labelled using custom software written in MATLAB R2021b (MathWorks, USA). To this end, signals from spheroids and brain slices were first denoised using Daubechies type 4 wavelet denoising. Spheroid signals were then also bandpass filtered using a 4^th^ order Butterworth filter; the high-pass frequency was set between 0.1 and 1 Hz, and was used to remove the baseline drifting likely due to the sub-optimal mechanical stability of the spheroid on the 3D MEA; the low-pass frequency was set at 300 Hz to remove multi-unit activity and isolate the field potential components. Automatic labels were always verified by visual inspection and manually corrected as required. EEG signals from humans were labelled manually by expert clinicians.

To compare the frequency response of ictal and interictal events in the four experimental groups (spheroid, brain slice, epileptic rat, epileptic human), we used the Cross Power Spectrum (CPS). To this end, all recordings were first filtered to eliminate powerline interferences with a band-stop filter between 48 and 52 Hz and a steepness of 0.99, and down-sampled to 2 kHz to reduce the computational demand of the analysis. Then, electroencephalographic relevant spectral bands were isolated with bandpass filters as follows: alpha band (8-12 Hz), beta band (15-30 Hz), gamma band (30-80 Hz), theta band (4-7 Hz), delta band (0.5-3 Hz). This band division was established to ease the comparison with human EEG and to enable the translation of the meaning of each result. The CPS was computed using the Welch’s method: the signals were divided in windows of 5 s with an overlap of 0.05 s; the Fast Fourier Transform of each window of the pair was multiplied and averaged to obtain an estimate of the CPS. The CPS threshold was set to 0, yielding positive values as an indicator of spectral features similarities. To eliminate the possibility of the high CPS values being a product of noise, we performed CPS against a negative control consisting of baseline noise, i.e., signal segments devoid of electrographic events, isolated from in vitro, in vivo and human recordings.

### Statistical Analysis

Statistical analysis was performed using Origin 8 (OriginaLab Corporation, USA) or SPSS Statistics 20 (IBM, USA). Data were first checked for normality and homoscedasticity to choose the appropriate test statistics. To compare the spheroid area size and maximum diameter from phase contrast images of the same samples across DIVs, we used one-way ANOVA for repeated measures, followed by the Tuckey post-hoc test. To compare the n. active/total spheroids, the occurrence of the three electrical phenotypes across DIVs, and the percentage of active electrodes we used the Pearson’s Chi-Square test accompanied by the Cramer’s V measure of association. In all cases, significance was set at p < 0.05. Data are expressed as mean ± SEM. Throughout the figures, we indicate the different p-value ranges as follows: * p < 0.05; ** p < 0.01; *** p < 0.001.

## Figures

Graphs were prepared in Microsoft Excel or GraphPad Prism; boxplots and electrophysiology signals were plotted in MATLAB R2021b. Figures were assembled in CorelDraw x8.

## Results

### High-yield and reproducibility of electrically active hippocampal spheroids

**Figure 1a** illustrates the protocol essential steps, consisting of the formation of cellular aggregates in U-shaped multiwell plates (step 1) and their subsequent transfer in agarose coated multiwell plates at day in vitro (DIV) 3 (step 2) to pursue the spheroid culture. **Figure 1b** shows phase contrast images of a representative spheroid throughout time. The spheroids increased their surface area (**Figure 1c**) and maximum diameter (**Figure 1d**) up to DIV21, and then stabilized at their final size (n = 15 spheroids/time-point, from three batches).

**Figure 1.**
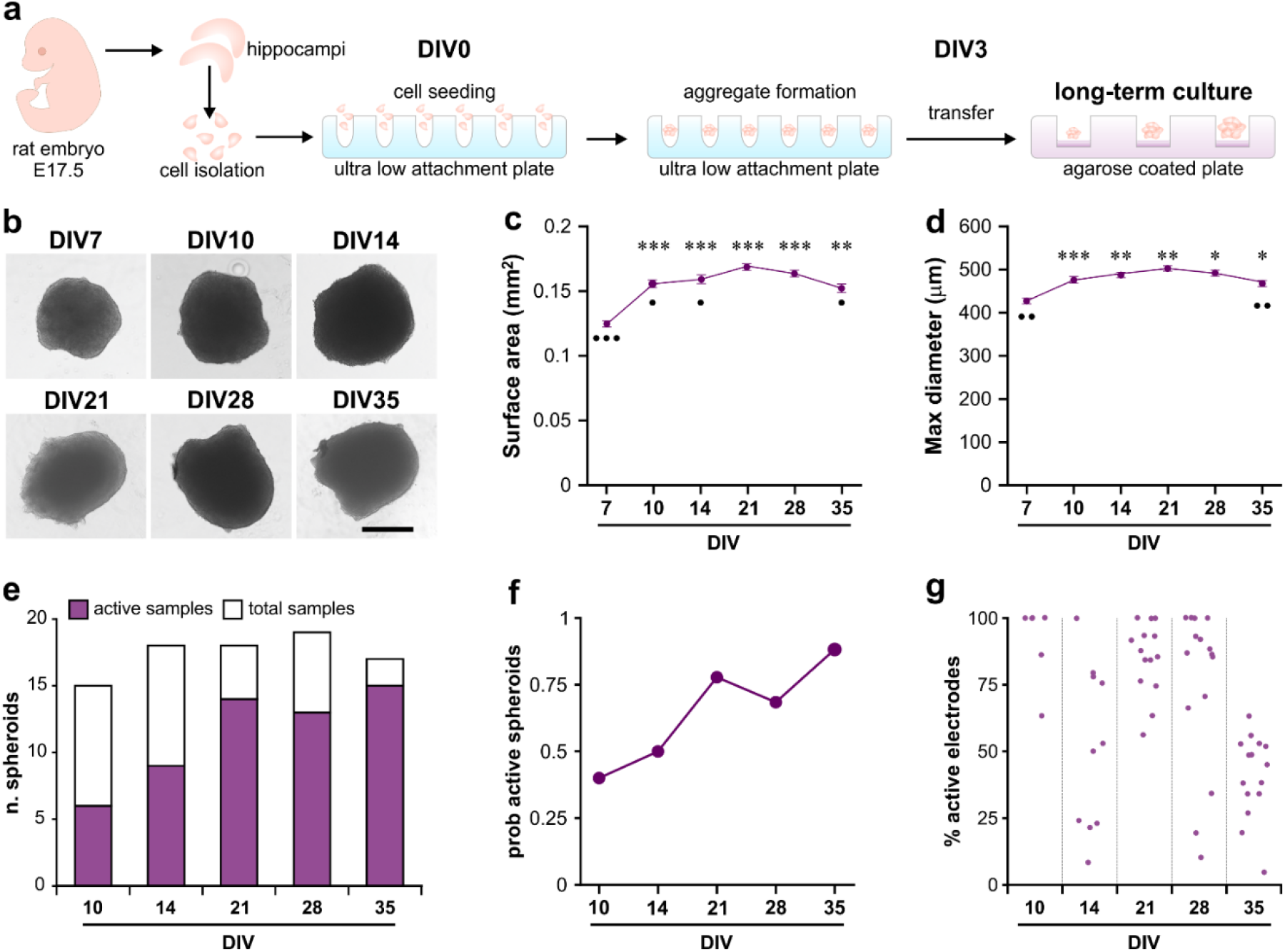
High yield of electrically active hippocampal spheroids for long-term electrophysiology studies. **a**. Schematic illustration of the two-step protocol for hippocampal spheroid static culture. Primary hippocampal cells are harvested from E17.5 rat embryos and seeded in ultra-low attachment multiwell plates to allow the formation of cellular aggregates. These are then transferred to an agarose-coated multiwell plate after 3 days in vitro (DIV) for long-term maintenance. **b**. Phase contrast microscopy images of spheroids across DIVs showing their increase in size with time. Scale bar: 250 µm. **c-d**. Results statistics for surface area (**c**) and maximum diameter (**d**) of the spheroids across DIVs indicate the achievement of their peak size at DIV21. One-way ANOVA repeated measures – area size, F(df): 32.78(5); maximum diameter, F(df): 12.34(5). *p < 0.05; **p < 0.01; ***p < 0.001 vs DIV7; **^•^**p < 0.05; **^••^**p < 0.01; **^•••^**p < 0.001 vs DIV21. **e**. Overview of the dataset of hippocampal spheroids acutely recorded with 3D microelectrode array. The superimposed bar charts of active and total spheroids indicate a high yield of electrically active spheroids across DIVs. **f**. The probability of finding electrically active spheroids (P_active_) increases with time. DIV10: P_active_ = 0.4; DIV14: P_active_ = 0.5; DIV21: P_active_ = 0.78; DIV28: P_active_ = 0.68; DIV35: P_active_ = 0.88. χ^2^(df): 11.39(4), p < 0.05; Cramer’s V = 0.36. **g**. Percentage of active electrodes observed with acute microelectrode array recordings across DIVs. Each dot represents an active spheroid. The maximum yield is obtained at DIV21-28.

To screen the electrical functionality of the spheroids over time, we performed acute electrophysiology recordings via 3D microelectrode array (MEA). To this end, individual spheroids were placed onto the MEA prior to the recording session and discarded afterwards. While acute MEA recordings do not portray the functional profile of the same spheroid in time (as opposed to chronic recordings), we preferred this approach to minimize the physical disturbances and mechanical constraint caused to the spheroid by the penetrating 3D electrodes; such disturbances may indeed affect the spheroid network self-organization and, in turn, influence its electrophysiological features. Overall, we screened n = 87 spheroids from 6 culture batches; of these, n = 60 spheroids were electrically active (global yield ∼70%). Electrical activity emerged at DIV10 and persisted until DIV35, i.e., the latest observation time-point (**Figure 1e**); further, the probability of finding electrically active spheroids increased with time (**Figure 1f**). The activity spread through the spheroid area, although the number of electrically active locations (‘active electrodes’) varied across the observed DIVs (**Figure 1g**). In particular, spheroids at DIV21-28 exhibited the most diffused activity, while at DIV35 the activity was confined to a smaller number of locations, suggesting that DIV35 should be the latest time point for reliable electrophysiology studies.

### Emergence of epileptiform patterns in hippocampal spheroids

As expected from developing neural networks, the spheroids exhibited different electrical phenotypes at different DIVs (**Figure 2a**). In this regard, we have serendipitously discovered that hippocampal spheroids exhibit spontaneous epileptogenesis. At early stages (DIV10-14), the spheroids generated short recurrent events consisting of an oscillatory pattern of population spikes overriding a slow field potential component (insert **(i)**). At the intermediate stage (DIV21), the spheroids started exhibiting a mixed pattern of short events (resembling those observed at DIV10-14, insert **(ii)**) and longer population bursts. The latter recurred as isolated events (insert **(iii)**) or in the form of barrages (insert **(iv)**), in which case they resembled the electrographic features of clonic ictal activity^21^. Burst barrages became the most robust and cohesive at DIV28-35, wherein they resembled full-blown ictal discharges with tonic-clonic features (insert **(v)**). Based on these observations, we will hereafter refer to these short and long electrographic entities as interictal-like and ictal-like events, respectively. As shown in **Figure 2b**, interictal-like events were initially longer in duration and recurred more slowly (DIV10-21); subsequently (DIV28-35), they became shorter and more frequent. On the contrary, ictal-like discharges were shorter and more frequent when they first emerged at DIV21, and reached their full-blown robustness at DIV28, when they also recurred at longer intervals (**Figure 2c**).

**Figure 2.**
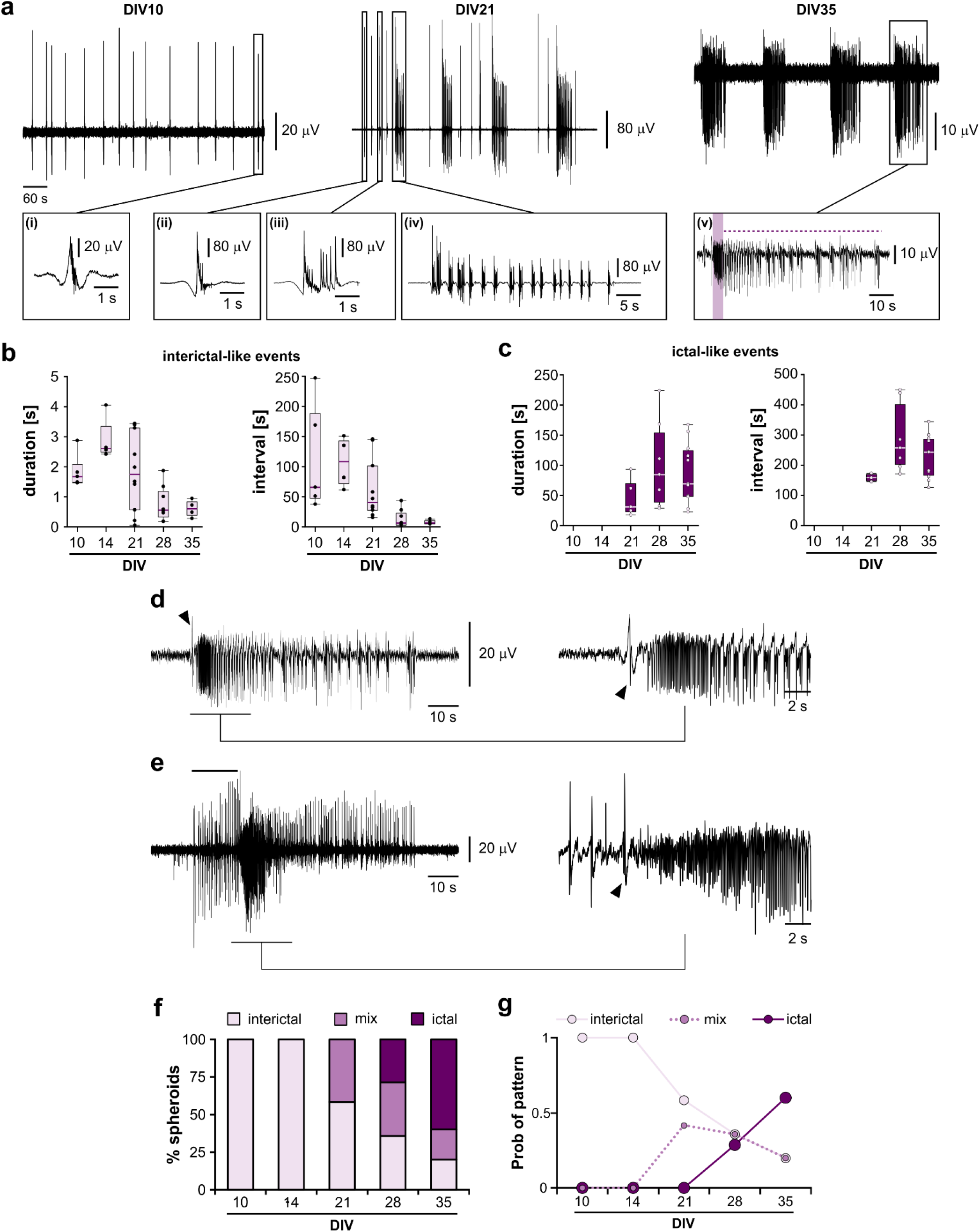
Hippocampal spheroids exhibit spontaneous epileptogenesis. **a**. Acute electrophysiology recording of hippocampal spheroids unveils their evolving functional features toward epileptogenesis. Representative signals from DIV10 (early stage), DIV21 (intermediate stage) and DIV35 (late stage). The boxed inserts show the indicated events at expanded time scales. At DIV10, the spheroids generated recurring short events consisting of population spikes overriding a slow field component – insert **(i)**. At DIV21, the spheroids started exhibiting a mixed pattern of short events – insert **(ii)**, long population bursts – insert **(iii)** and burst barrages – insert **(iv)** resembling the electrographic features of clonic ictal activity. At DIV35, the spheroids started generating full-blown ictal-like activity with tonic and clonic features, indicated in insert **(v)** by the purple shading and the dashed purple line, respectively. **b**. Box plots summarizing the duration and interval of occurrence of interictal-like events across DIVs show a trend toward shorter and more frequent discharges with time. Horizontal lines indicate the median values, whiskers are the minimum and maximum values, each dot is a sample. **c**. Box plots summarizing the duration and interval of occurrence of ictal-like events across DIVs show their emergence at DIV21 and a trend toward longer duration and interval of occurrence with time. Horizontal lines indicate the median values, whiskers are the minimum and maximum values, each dot is a sample. **d-e**. Unfolding of ictogenesis in hippocampal spheroids. In **(d)** is a representative tonic-clonic ictal discharge riding abruptly over an interictal-like event (arrowhead); in **(e)** is a representative tonic-clonic ictal discharge announced by a barrage of interictal-like events (solid line). The inserts on the right of each representative trace show their respective ictal onset at expanded time scale, where the arrowhead points to the initiating interictal discharge. **f**. The electrical phenotype of hippocampal spheroids changes in time. The stacked column graph shows that early-stage spheroids (DIV10-14) exhibit a purely interictal-like pattern, which is more rarely encountered at later DIVs, with the majority of spheroids generating ictal-like activity at DIV35. **g**. The probability of observing the three electrical phenotypes is associated with the spheroid DIV (χ^2^(df): 29.29(8), p < 0.001; Cramer’s V = 0.52).

Ictal-like activity exhibited clonic-like features with abrupt onset in n = 2 spheroids only (*cf*. **Figure 2a** (iv)), while the vast majority of the events exhibited tonic-clonic components. The latter were always initiated by an interictal-like discharge; in 66.7% of the spheroids, they emerged abruptly with a tonic-like oscillatory pattern immediately following or riding over the initiating interictal-like event (**Figure 2d**), whereas in 33.3% of the cases they were announced by a barrage of interictal discharges (**Figure 2e**). In our hands, the two types of pre-ictal state never coexisted, suggesting unique ictogenic mechanisms and their spheroid-specificity.

The probability of observing each of the three patterns (interictal-like only, predominantly ictal-like, mixed) followed the time of the culture (**Figure 2f-g**). Specifically, spheroids at DIV10-14 generated interictal activity only, whereas at later stages they could generate any of the three patterns with different probabilities across DIVs. This feature suggests the developmental unfolding of epileptogenesis as well as the peculiarity of each spheroid in terms of the generated epileptiform pattern.

### Epileptiform patterns generated by hippocampal spheroids resemble those of animal epilepsy models and of epileptic humans

To corroborate the pathophysiological identity of the electrographic events generated by the spheroids, we compared their spectral features against those of known interictal and ictal events recorded from the hippocampus of established in vitro and in vivo rodent models, as well as of drug-resistant epileptic humans. As in vitro model we used mouse hippocampus-cortex slices treated with 4-aminopyridine^22^; as in vivo model, we used kainic acid-treated epileptic rats^20^. **Figure 3a** and **b** show representative ictal and interictal discharges, respectively, and their spectra. Along with the electrographic similarities, the spectra suggested the presence of similar frequency components within interictal and ictal events recorded from the animal models and the human epileptic hippocampus. To validate this observation, we performed cross power spectrum (CPS) analysis. The CPS yields the cross-correlation between two signals in their frequency domain; thus, it provides information about the frequency components shared by the two signals, wherein positive CPS values indicate common frequencies. As shown in **Figure 3c-d**, the CPS confirmed the presence of common frequency components, yielding the strongest values in the delta (δ, 0.5-3 Hz) and theta (θ, 4-7 Hz) frequency ranges for both event types. Remarkably, the CPS peak values of ictal discharges concur with what observed in MTLE patients with hippocampal sclerosis^23^. As expected, no similarity was found between the spheroid signals and baseline segments devoid of electrographic events (**Figure 3e**), confirming that the positive CPS values are not a product of noise, but reflect true common spectral features.

**Figure 3.**
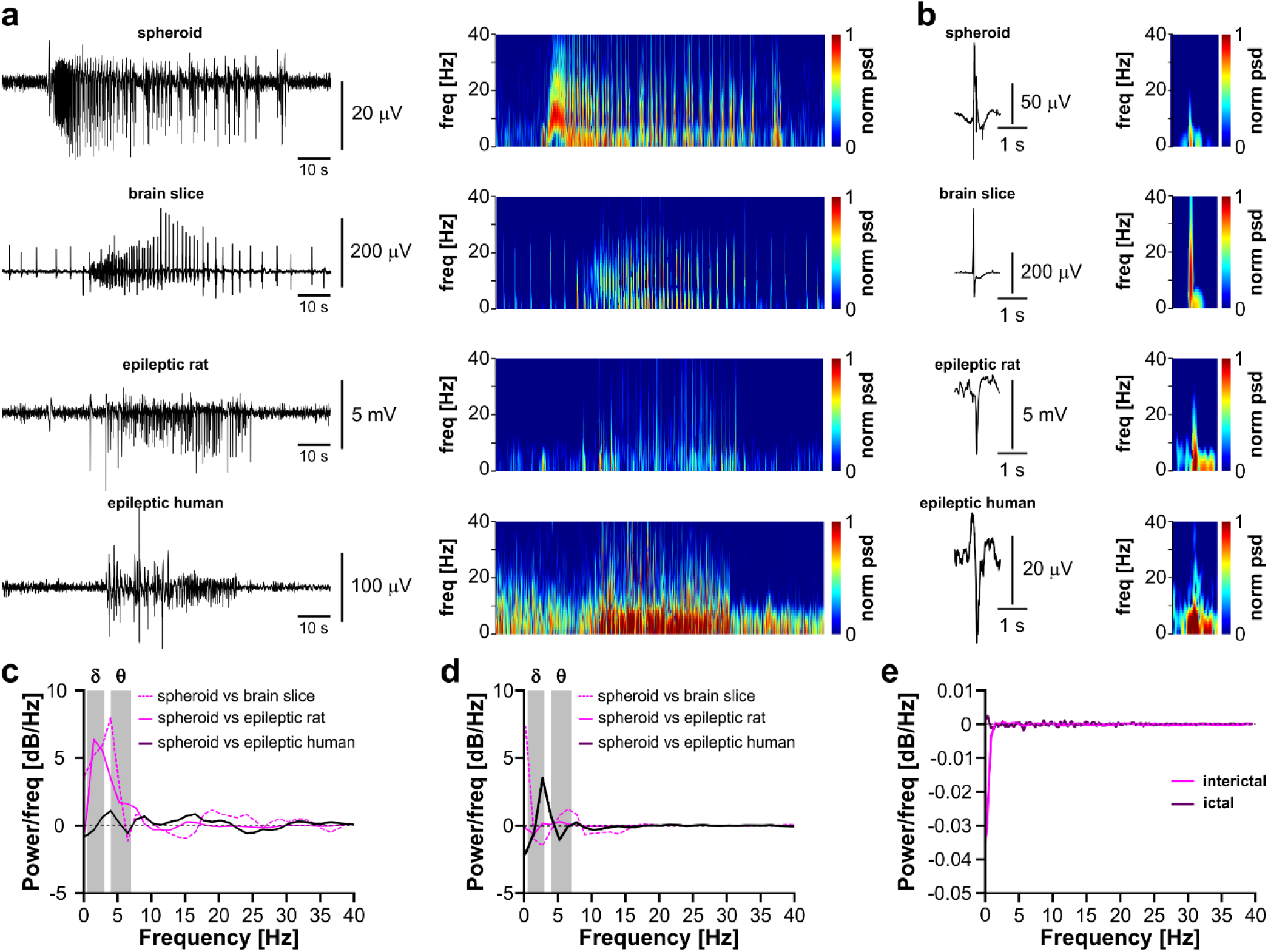
Epileptiform patterns generated by hippocampal spheroids recapitulate the spectral features of in vitro and in vivo animal models as well as of human epilepsy. **a**. On the left are representative ictal discharges recorded from hippocampal spheroids, and the hippocampus of a 4-minopyridine-treated hippocampus-cortex slice, a kainic acid-treated epileptic rodent and a drug-resistant epileptic human. At the right of each recording is the respective spectrum. In the brain slice spectrum, the white arrow points to the onset of the ictal discharge superimposed on the CA3-driven interictal events, inherent to this model. **b**. On the left are representative interictal discharges recorded from hippocampal spheroids, and the hippocampus of a 4-aminopyridine-treated hippocampus-cortex slice, a kainic acid-treated epileptic rodent and a drug-resistant epileptic human. At the right of each recording is the respective spectrum. In **a-b** the color bar spans a power spectral density (psd) range of 25 db/Hz; because of the different thresholds required across signals, the color bar is normalized in the range [0 1] for easier interpretation, where 0 and 1 are the minimum and maximum psd, respectively. **c-d**. Cross power spectrum (CPS) of ictal **(c)** and interictal events **(c)** generated by the spheroids against known events recorded from established animal models and epileptic humans. The strongest CPS values are in the range of the delta (δ, 0.5 to 3 Hz) and theta (θ, 4-7 Hz) bands, both for ictal **(c)** and interictal **(d)** events. For the latter, the high CPS values at very low frequencies in the comparison with brain slice signals are to be considered an artifact due to the short nature of interictal events. **e**. No positive CPS values are detected when comparing the spheroid signals with the isolated baseline segments (negative control), confirming that the CPS values are not a product of noise.

## Discussion

The dysfunction and aberrant rewiring of the hippocampus play a primary role in MTLE, the most common form of epilepsy in adults. Given the pivotal involvement of the hippocampus in cognitive and emotional processes, its functional compromise in MTLE not only contributes to seizure manifestation; it is also linked to the cognitive and psychiatric co-morbidities often seen in MTLE patients. Along these lines, functional and structural hippocampal alterations are also observed in patients with dementia. While medical therapy is only symptomatic and cannot cure the condition, regenerative medicine promises to overcome this limitation. However, brain regeneration is still in its infancy and as such pre-clinical safety studies are still needed. These are of utmost importance for devising effective and reliable regenerative strategies and prevent iatrogenic worsening of the treated condition. Here, we have characterized the electrophysiological behavior of primary hippocampal spheroids cultured in physiological medium. Our findings demonstrate the intrinsic epileptogenicity of hippocampal cells when they are allowed to self-organize in 3D assemblies.

Previous work has shown that 3D bioengineered hippocampal networks relying on silica microbead scaffolds exhibit a more robust electrical activity than their 2D counterpart^24^; however, the generation of ictal discharges was never observed either spontaneously or upon electrical stimulation. Spontaneous epileptiform discharges in cultured hippocampal neurons were only reported in 2D microcultures in the early 1990’s^12,13^, and they were ascribed to the formation of recurrent excitatory connections and autapses consequent to the ultra-low culture density. Overall, this evidence suggests that the hyper-excitable phenotype of the hippocampal spheroids described here may result from their self-organization in the absence of mechanical constraints. To the best of our knowledge, our work brings the first evidence of such epileptogenic propensity.

Other studies have addressed epileptiform network dynamics using stem-cell derived organoids, but they had to rely on pharmacological manipulation^25^, or on the combination of targeted genetic mutation and pharmacological treatment^26^ to attain the pathological phenotype. While brain organoids have become a fashionable tool to study developmental epilepsies, they still suffer for a high intrinsic variability^27^ and they are inherently bound to specific genetic mutations linked to epilepsy; the latter does not permit to study epileptic syndromes of unknown origin, which make up more than one half of the epilepsies. Further, stem cell-derived brain organoids entail high costs and long culturing time (months) to achieve a yet functionally immature phenotype. The robust network activity displayed by hippocampal spheroids manifests the importance of using primary cells over embryonic stem cells, neural progenitors, or induced pluripotent stem cells, as these hardly achieve spontaneous electrophysiological function, despite the expression of synaptic components, ion channels and neurotransmitters demonstrated by transcriptomic analysis. For these reasons, primary brain cells are re-gaining attention for producing spheroids thanks to the faster maturation, high reproducibility and moderate costs^28^. These features support reliable high-throughput platforms to address neuronal function in health and disease as well as to study novel treatments for brain disorders.

A key aspect of epilepsy is the re-organization of brain circuitry, wherein seizures represent aberrant network phenomena. However, current 3D models are presented with a deep characterization of cellular phenotypes and molecular signatures, whereas information on the global network dynamics remains elusive. In fact, most of the functional characterization studies have relied on whole-cell patch clamp and single/multi-unit electrophysiology, and on calcium imaging^25,26^, which resolve network activity at the single cell or small cell ensemble level; these techniques fail to provide a high-level portrait of the network dynamics typical of epilepsy, and as such they do not support a direct comparison with what observed in the EEG of epileptic patients. At variance, field potential recording by means of 3D MEA has enabled us to address the global network dynamics of hippocampal spheroids and unveil their epileptiform behavior.

The value of primary neural spheroids for high-throughput basic and translational research is also denoted by the spontaneous emergence of oscillatory activity in spheroids obtained from rodent primary cortical cells, reminiscent of synchronous network oscillations seen in vivo^29^. Our work corroborates this translational value by showing an unprecedented parallelism of epileptiform patterns spontaneously generated by hippocampal spheroids with the electrographic and spectral features of those generated by the human epileptic hippocampus.

## Conclusions and Future Directions

We have demonstrated the inherent emergence of epileptiform patterns in developing hippocampal spheroids. The probabilistic occurrence of any of the interictal, ictal or mixed electrical phenotypes in the absence of a known epileptogenic insult brings an important parallelism with human acquired epilepsies of unknown etiology. These findings warrant caution in devising hippocampal cell-based therapies for regenerative medicine, as they may ensue epileptogenesis. In this regard, it would be highly relevant to evaluate the functional features of hippocampal spheroids when integrated with other brain structures (e.g., cortex) to form assembloids mimicking the complexity of the hippocampus-cortex loop. Further, in vivo studies complementing this and previous work^11,15^ will shed more light into the behavior of hippocampal grafts in intact animals. To this end, chronic non-invasive electrophysiological recordings, e.g., through embedded stretchable mesh nanoelectronics^30^, will enable the deployment of machine learning techniques to forecast the emergence of pathological electrical activity, thus supporting pre-screening of tissue replicas for safe brain regeneration. At the same time, hippocampal spheroids cultured and recorded in physiological medium are an invaluable tool to address the mechanisms of epileptogenesis and ictogenesis in a simplified and high-throughput model. In turn, this will enable their effective application as test-bed in the design and screening of new therapeutic approaches for epileptic syndromes involving the hippocampus.

## Author Contributions

**John W. Ephraim:** Conceptualization, Methodology, Validation, Investigation, Formal analysis, Data curation, Visualization, Writing - Original Draft, Writing - Review & Editing. **Davide Caron:** Investigation, Formal analysis, Data curation, Visualization, Writing - Review & Editing. **Angel Canal-Alonso:** Formal analysis, Data curation, Visualization, Writing - Original Draft, Writing - Review & Editing. **Juan Manuel Corchado:** Resources, Formal analysis, Data curation, Supervision, Writing - Review & Editing, Funding acquisition, Resources, Project administration. **Gemma Palazzolo:** Conceptualization, Methodology, Validation, Investigation, Formal analysis, Data curation, Visualization, Supervision, Writing - Original Draft, Writing - Review & Editing, Funding acquisition, Project administration. **Gabriella Panuccio:** Conceptualization, Validation, Formal analysis, Data curation, Visualization, Supervision, Writing - Original Draft, Writing - Review & Editing, Funding acquisition, Resources, Project administration.

## Acknowledgements

This work was funded by the European Union under the Horizon 2020 work program, FET-PROACT(RIA), project HERMES – Hybrid Enhanced Regenerative Medicine Systems, Grant Agreement n. 824164.

